# Development of a floxed *Gabbr2* gene allows for widespread conditional disruption of GABBR2 and recapitulates the phenotype of germline Gabbr2 knockout mice

**DOI:** 10.1101/2025.01.23.634473

**Authors:** Julie R. Hens, Stacey Brown, Pawel Licznerski, Jacqueline Suarez, Elizabeth Jonas, John J. Wysolmerski

## Abstract

GABBR1 and GABBR2 are widely expressed in the brain and genetic inhibition of their function leads to widespread neurologic dysfunction and premature death in mice. Given that GABBR1 and GABBR2 heterodimerize to form a functional receptor, global knockout of GABBR1 or GABBR2 results in a similar phenotype, characterized by spontaneous epileptiform activity, hyperlocomotor activity, hyperalgesia, impaired memory and premature death. It is now known that both GABBR1 and GABBR2 are expressed in a variety of tissues outside the nervous system and that GABA-B receptors can heterodimerize with other class C GPCRs, including the extracellular calcium-sensing receptor (CaSR). Studies in vitro have demonstrated that interactions with GABBR1 and GABBR2 can alter CaSR signaling in human embryonic kidney cells and breast cancer cells. The neurologic consequences of global loss of function of GABBR1 or GABBR2 has made it difficult to study the effects of loss of GABBR function in other organs. While a conditional knockout for GABBR1 is available, the GABBR2 gene had not been “floxed”. We have used CRISPR to insert loxP sites into the GABBR2 locus in mice. These mice are normal at baseline but when bred with mice expressing Cre-recombinase under the control of the ubiquitously expressed Actin gene promoter, they recapitulate the phenotype of global GABBR2 knockout mice. Phenotypic changes through the brain, including the cortex, hippocampus and cerebellum. Evidence of abnormal neuronal function, increase cell death, and changes in neuronal architecture are seen throughout the brain of CRISPR knockout mice. These mice should be useful tools to study cell type-specific loss of GABBR2 function in the brain and other organs.

## INTRODUCTION

The GABA-B receptors (GABBR1 and GABBR2) are class C, G-protein-coupled receptors (GPCRs) that heterodimerize to form a receptor complex responding to gamma-aminobutyric acid (gaba), the major inhibitory neurotransmitter in the brain. The GABBR1 subunit contains the gaba-binding site, whereas the GABBR2 subunit is responsible for interacting with G-proteins. Furthermore, GABBR1 contains an endoplasmic reticulum (ER) retention site, which prevents its trafficking to the plasma membrane. However, heterodimerization with GABBR2 allows interactions between the coiled-coil sequences of each subunit, masking the ER retention site in GABBR1, and allowing translocation of the heterodimeric complex to the plasma membrane. Most commonly, the heterodimeric receptor couples to G_i_ or G_o_, leading to inhibition of adenylate cyclase activity, inositol triphosphate synthesis, voltage-gated calcium channels, and potassium channels (1, 2). As a result, GABABRs hyperpolarize neurons and inhibit the release of several neurotransmitters, resulting in the suppression of neuronal activity in many brain areas.

GABBR1 and GABBR2 are widely expressed in the brain and genetic inhibition of their function leads to widespread neurologic dysfunction and premature death in mice. Given that GABBR1 and GABBR2 heterodimerize to form a functional receptor, global knockout of GABBR1 or GABBR2 results in a similar phenotype, characterized by spontaneous epileptiform activity, hyperlocomotor activity, hyperalgesia, impaired memory and premature death (3). As these results demonstrate, GABA-B receptors clearly have important functions in the brain. However, it is now known that both GABBR1 and GABBR2 are expressed in a variety of tissues outside the nervous system (3–6). Furthermore, it has been shown that the GABA-B receptors can heterodimerize with other class C GPCRs, including the extracellular calcium-sensing receptor (CaSR)(7, 8). Studies *in vitro* have demonstrated that interactions with GABBR1 and GABBR2 can alter CaSR signaling in human embryonic kidney (HEK) cells and breast cancer cells (5, 9). Furthermore, the CASR and GABBR1 interact in chondrocytes in the growth plate and in parathyroid cells *in vivo* (4). Recent studies have demonstrated that heterodimerization of GABBR1 and the CaSR in the parathyroid glands modulates calcium-mediated PTH secretion and systemic calcium metabolism (10), demonstrating that GABBR’s can regulate signaling from other receptors.

The neurologic consequences of global loss of function of GABBR1 or GABBR2 has made it difficult to study the effects of loss of GABBR function in other organs. While a conditional knockout for GABBR1 is available, the GABBR2 gene had not been “floxed”. Therefore, to study the interactions between the CaSR and GABBR2 in organs other than the brain, we have used gene editing techniques to insert loxP sites into the GABBR2 locus in mice. These mice are normal at baseline but when crossed with mice expressing Cre-recombinase under the control of the ubiquitously expressed Actin gene promoter, they recapitulate the phenotype of global GABBR2 knockout mice. These mice should be useful tools to study cell type-specific loss of GABBR2 function in the brain and other organs.

## METHODS

### Generation and breeding of Gabbr2 cKO Mice

The GABBR2 cKO mouse model was generated via CRISPR-Cas9 genome editing (11, 12) (13). Potential Cas9 target guide (protospacer) sequences in introns 9 and 10 were screened using the online tool CRISPOR http://crispor.tefor.net (14) and candidates were selected. Templates for sgRNA synthesis were generated by PCR, sgRNAs were transcribed *in vitro* and purified (Megashortscript, MegaClear; ThermoFisher). sgRNA/Cas9 RNPs were complexed and tested for activity by zygote electroporation, incubation of embryos to blastocyst stage, and genotype scoring of indel creation at the target sites. The sgRNAs that demonstrated the highest activity were selected for creating the floxed allele. Guide RNA (gRNA) sequences are as follows: intron 9, 5’ guide: ACTAGATCCTCTCACCCAGT and intron 10, 3’ guide CTGCCATGCTGTGACCCCAT. Accordingly, a 615 base long single-stranded DNA (lssDNA) recombination template incorporating the 5’ and 3’ loxP sites was synthesized (IDT). The C57Bl6 3 SJL F2 or FVB/NJ zygote embryos were transferred to the oviducts of pseudopregnant CD-1 foster females using standard techniques(13, 15). Genotype screening of tissue biopsies from founder pups was performed by PCR amplification and Sanger sequencing to verify the floxed allele. Germline transmission of the correctly targeted allele (i.e., both loxP sites *in cis*) was confirmed by breeding and sequence analysis. Seven potential founders with a floxed *Gabbr2* gene were identified, and three (#33, #14, #19) true-breeding FVB lines were generated. We also generated two true-breeding lines on a C57bl/6 mouse background. The studies described herein were performed on animals derived from lines 33 and 19. The two lines were maintained separately, but because of their similar biochemical phenotypes, data from the two lines have been pooled except where indicated.

We crossed *GabbR2* ^lox/lox^ mice with B6.FVB-*Tmem163^Tg(ACTB-cre)2Mrt^*/EmsJ (Actin-cre) to verify the effectiveness of the CRISPR-generated lox sites on *GabbR2*. The resulting offspring were then crossed again to *GABBR2* ^lox/lox^ mice (control) to generate Actin-Cre/*GABBR2* ^lox/lox^ mice (cKO).

All studies described in this manuscript were performed on animals between the ages of 3 and 15 weeks unless specifically indicated. All procedures were per Yale University Animal Care and Use Committee and U.S. National Institutes of Health standards.

### RNA and protein analysis

Brains from 3-week-old mice were removed and total RNA was isolated. One μg of RNA was converted to cDNA using Applied Biosystems high-capacity cDNA reverse transcription kit (Thermo Fisher Scientific, Waltham, MA). Taqman probes were used to measure GABBR1(Mm00444578_m1), GABBR2(Mm01352554_m1), CASR (Mm00443375_m1), and GAPDH (#4352339E) (Thermo Fisher Scientific, Waltham, MA). Real-time PCR was performed using TaqMan ^TM^ Fast Universal PCR Master Mix reagents (Thermo Fisher Scientific, Waltham, MA) and Applied Biosystems StepOne Plus Real-Time PCR System. Ct values were analyzed using the ΔΔ – Ct method (16).

For protein isolation, half the brain cut in the coronal mid-line was added to 1 ml of RIPA buffer (50 mM Tris HCl pH 8, 150 mM NaCl, 1% NP-40, 0.5% sodium deoxycholate, 0.1% SDS) with complete mini protease inhibitors (Roche Diagnostics, Mannheim, Germany). Using a TissueLyser II (Qiagen, Germantown, MD) with a 5 mm bead, tissue was lysed for 2 minutes at 30 rotations per second. Lysates were incubated on ice for an hour, before being centrifuged at 12,000 g, for 20 minutes. Thirty micrograms of protein were loaded in a well. Samples were not heated, and after the transfer, blots were blocked for 1 hour in 5% milk with 0.1 % Tween-20. Primary antibodies were added at 1/1000 overnight at 4^0^C while rocking. Blots were washed with PBS, and then goat-anti rabbit or goat-anti mouse secondary antibody was added for an hour, samples were washed in PBS. Blots were imaged using Odyssey Li-Cor system. Results were normalized to actin.

We used antibodies to Gabbr1 (#ab55051, Abcam, Waltham, MA), Gabbr2 (#ab181736, Abcam, Waltham, MA), Casr (#ACR-004, Alomone, Limerick, PA), actin (#MA5-11869, Invitrogen, Rockford Illinois), IRDye® 800CW Goat anti-Mouse IgG (H + L) (Li-Cor, Lincoln, Nebraska) IRDye®, 680RD Goat anti-Rabbit IgG Secondary Antibody (Li-Cor, Lincoln, Nebraska)

### Histology

Brains were paraffin-embedded and 5-micron sections were acquired. Sections were stained with hematoxylin and eosin, Luxol fast (17), or immunohistochemistry was performed with S100 antibody to examine changes in myelination in the central nervous system. Embedding and staining of mice brain tissue was done through Yale Pathology Tissue services.

### Motor agility

To examine motor changes in cKO compared to control mice, we assessed rotarod performance. Mice were trained to stay on the rotarod (AccuScan Instruments) (12 rpm) for 300 sec over two separate sessions the day before the experiment. During the test day, the length of time each mouse remained on the cylinder (“endurance time”; a maximal score of 300 sec) was measured immediately before (time 0) and 1, 2, and 4 hours after the application of L-baclofen (12.5 mg/kg) or vehicle (saline). The dose of baclofen that showed maximal effects on rotarod performance was determined in previous studies (3, 18).

### Behavioral experiments

We examined hyperactivity and unsupported rearing to analyze activity in the mice. Male cKO mice between 6 and 8 weeks of age were used for all experiments. Before behavioral testing, the investigator individually handled mice (3 times over 72 hours before the test day) to decrease anxiety. Next, mice were placed in a new, empty home cage where unsupported rearing and locomotor activity were monitored for 10-minute sessions, video recorded, and the last 5 minutes were scored manually. Unsupported rearing was defined as rearing without any contact with the walls of the test cage. The investigator was blinded as to the genetic variant during scoring.

### Statistical analysis

Data are presented as mean± standard error (SE). Comparisons between two groups were conducted using Student’s unpaired two-tailed t-tests. Where appropriate, two-way ANOVA with Sidak multiple comparison tests were used. All analyses were performed using Prism 10 (GraphPad Software, La Jolla, CA).

## RESULTS

### Insertion of LoxP sites and reduction in GABBR2 expression

Using CRISPR we inserted loxP sites into 5’ and 3’ sites flanking exon 10 of the *Gabbr2 gene*, which encodes the first transmembrane domain of the receptor (Figure 1a). We targeted this exon for several reasons. First, it was predicted to result in the loss of the first transmembrane domain. Second excision of this portion of DNA was predicted to result in a frameshift and mistranslation of all downstream exons when the primary transcript was spliced. Both of these characteristics are likely to result in a nonfunctional protein that would be degraded. Finally, targeting this relatively small exon allowed both flanking loxP sites to be targeted with one oligomer, allowing for more efficient editing.

**Figure 1.**
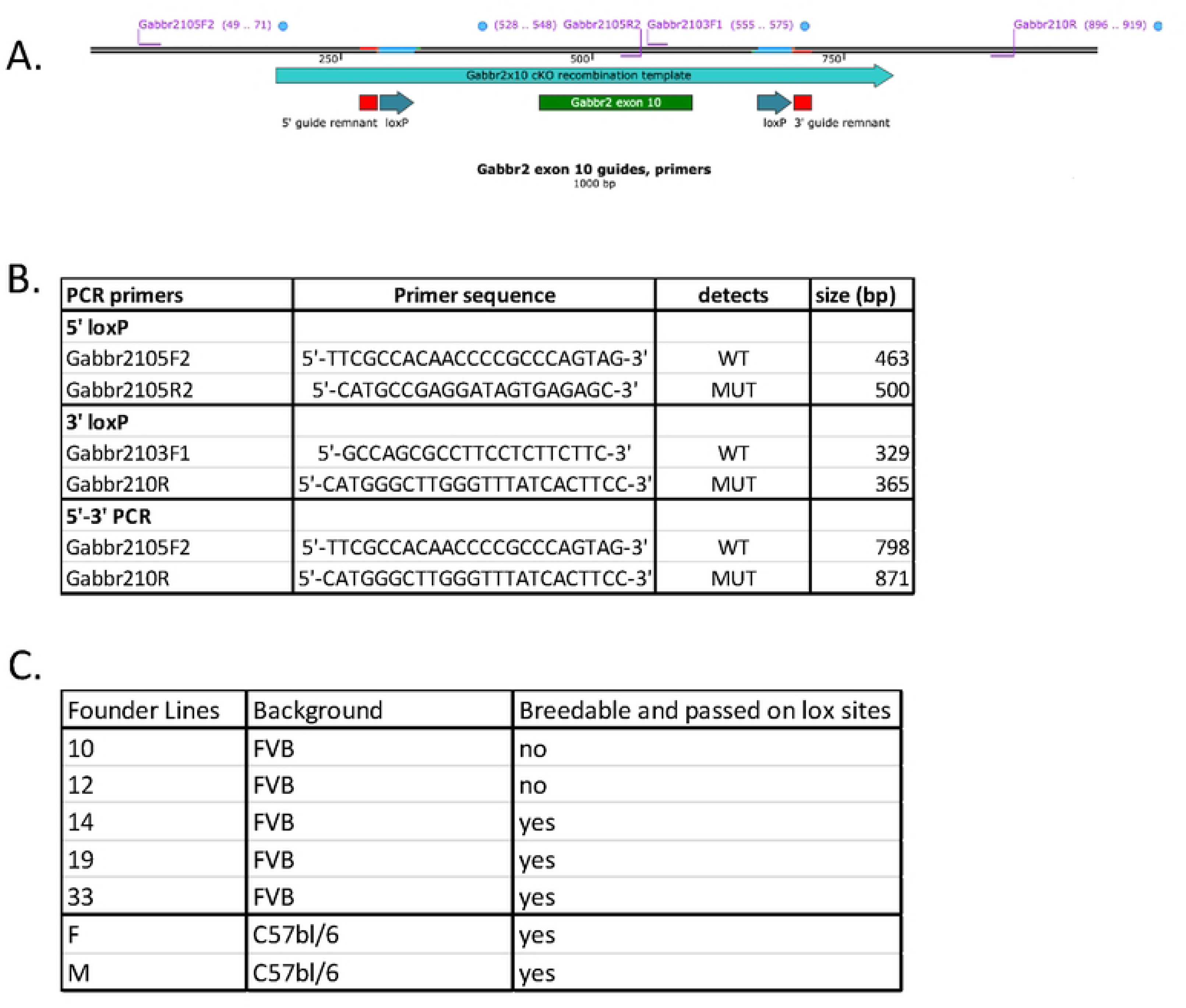
CRISPR design to add loxP sites to the *Gabbr2* gene. A. Map of *Gabbr2* gene showing the guide RNAs used to create LoxP sites. B. Primers used to identify loxP sites in Actin-CRE Gabbr2^lox/lox^ mice. C. Table summarizing the different Gabbr2^lox/lox^ mouse lines generated.

Primers were designed to detect wild-type and loxP sites at the 5’ and 3’ end of exon 10 to detect the appropriately floxed alleles (Figure 1b). Using these primers, we identified 5 potential founder lines in a FVB background that contained both loxP sites, three of which passed on the correct allele in a Mendelian fashion. We also identified 2 founder lines in a C57Bl/6 background, both of which passed on the correct genotype to offspring in Mendelian fashion (Figure 1c). We used lines 19 and 33 in an FVB background, (referred to as Gabbr2^lox/lox^ mice) in the following experiments.

Gabbr2^lox/lox^ mice were bred to B6.FVB-*Tmem163^Tg(ACTB-cre)2Mrt^*/EmsJ (Actin-cre) mice to generate Gabbr2 cKO mice with widespread loss of GABBR2 expression. In order to verify the loss of GABBR2, we examined *Gabbr2 mRNA* levels in whole brains from 21-day-old mice. *Gabbr2* mRNA expression was reduced by 80% in the Gabbr2 cKO mice as compared to Gabbr2^lox/lox^ (control) mice, lacking Cre expression. Loss of *Gabbr2 mRNA* expression did not affect either *Gabbr1mRNA* or *Casr mRNA* levels, two potential heterodimerization partners for GABBR2 (Figure 2A). We assessed GABBR2 expression by immunoblots of whole brain extracts. As shown in Fig. 2B, no GABBR2 protein was detected in extracts of whole brains harvested from cKO mice, although it was easily detected in brain extracts from control mice. As with the mRNA levels, loss of GABBR2 protein did not affect GABBR1 or CASR protein levels (Fig. 2B). These results demonstrate the effective elimination of GABBR2 expression when Gabbr2^lox/lox^ mice are bred with Cre recombinase-expressing mice.

**Figure 2.**
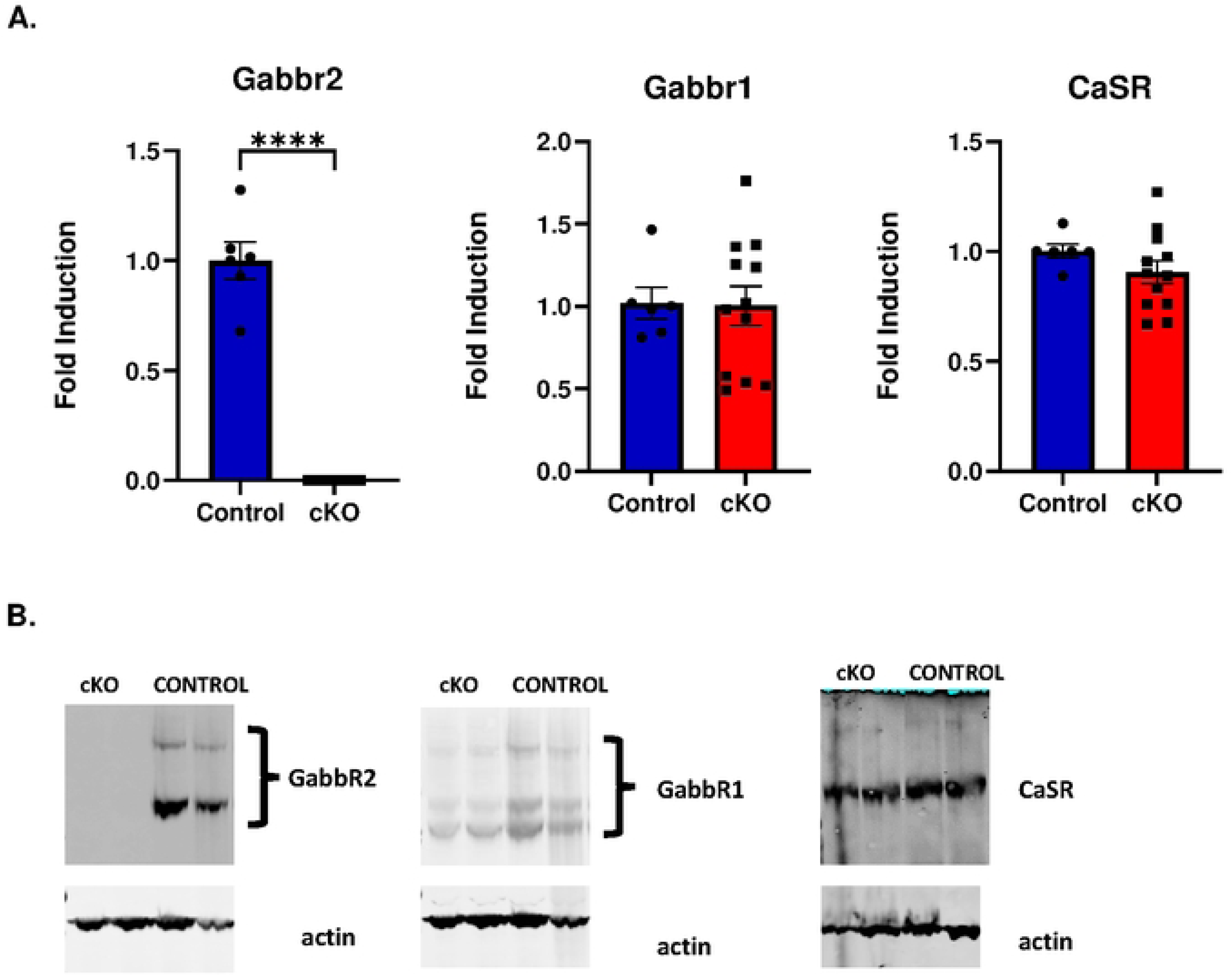
Expression of *Gabbr2*, *Gabbr1*, and *Casr* in the brains of control and cKO mice. A. QPCR to assess mRNA levels in whole-brain RNA. The specific transcript is shown on the top of each graph. Bars represent the mean ± SEM. *** p<.001. B. Protein expression levels of GABBR2, GABBR1, and CASR in whole-brain extracts. n= 6 control and n=12 cKO.

### Histological changes in the brain due to the loss of GABBR2

GABBR2 is expressed throughout the brain, including the cerebral cortex, cerebellum, Purkinje neurons, hippocampus, CA3 neurons, thalamic nuclei, medial habenula, and astrocytes (19–23). S100 proteins are expressed diffusely in glial cells, astrocytes and neurons throughout the brain (24, 25). In the GABBR2 cKO cortex, there was a generalized decrease in diffuse S100 staining and fewer distinct S100-positive cells when compared to control mice (Figure 3A versus 3B, red arrows). In addition, there were fewer S100-positive dendritic extensions in the GABBR2 cKO cortex (Figure 3A versus 3B, yellow arrows). There was also an increase in vacuolated neuronal bodies and cell debris evident in Luxol blue stained sections (Figure 3C versus 3D, green arrows), suggesting potential neuronal damage.

**Figure 3.**
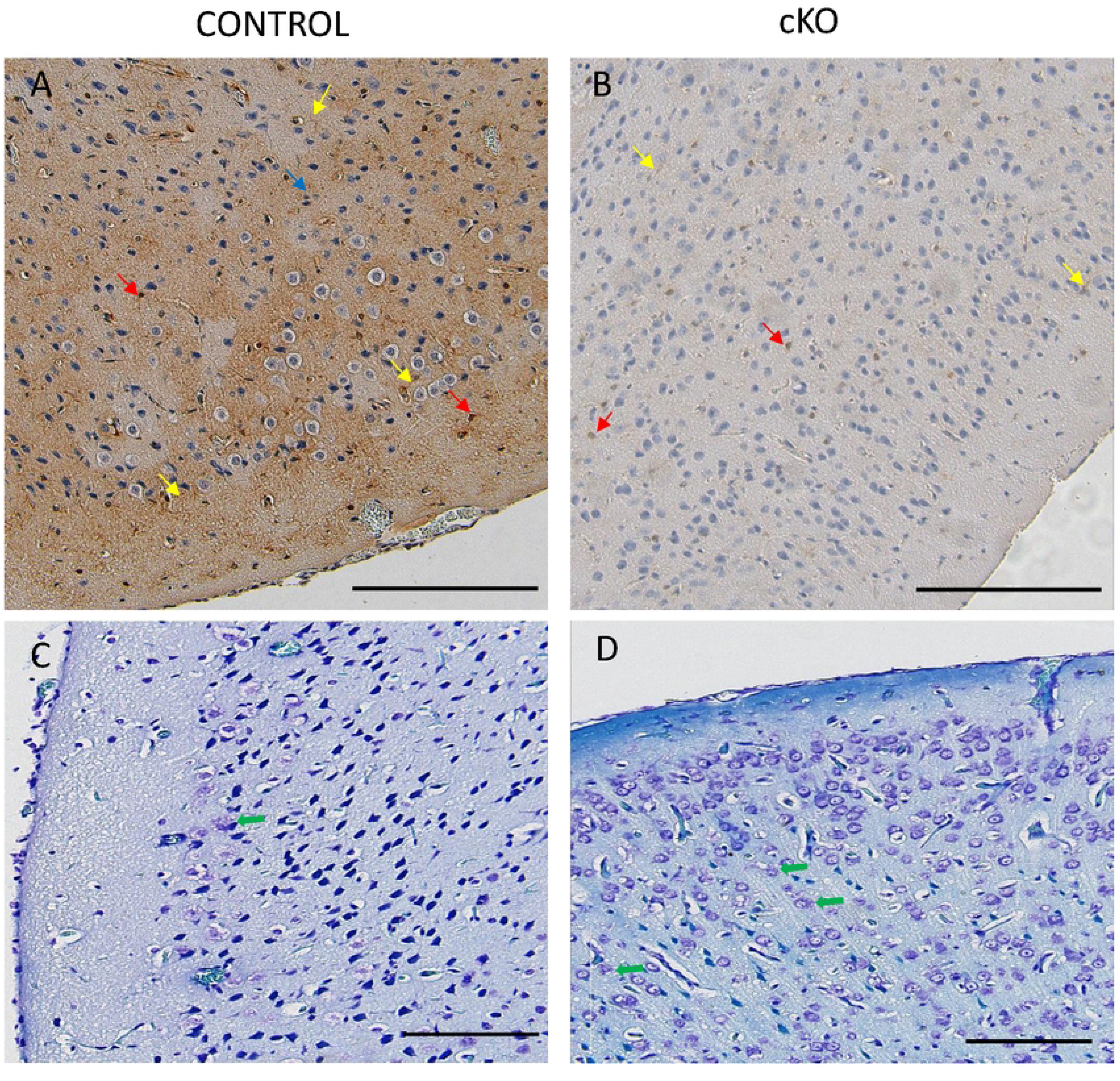
Brain sections of cerebral cortex from cKO and control brains stained for S100 and Luxol blue. There was a reduction of S100 staining overall and fewer S100-labeled neurons in the cKO cortex (A) versus control cortex (B). Yellow arrows point to neuronal dendrites. Red arrows point to S100-labeled neuronal bodies. There are more vacuolated neurons and cell debris in the cortex of cKO mice revealed by Luxol blue staining (C versus D). Green arrows point examples of vacuolated neurons. Scale bar = 200 microns.

Changes were also evident in the dentate gyrus and CA3 region of the hippocampus of GABBR2 cKO mice. There was a clear reduction in staining of the CA3 region (Figure 4A versus 4B, blue arrows). We observed a clear reduction in the number and layers of dense immature granular cells (Figure 4A versus 4B, green arrows and dotted border) as well as an increase in vacuolated cytoplasm in granular cells (Figure 4A versus 4B, and Figure 4C versus 4D, red arrows). There were many shrunken pyramidal cells in the polymorphic cell layer (Figure 4A and 4B, yellow arrows).

**Figure 4.**
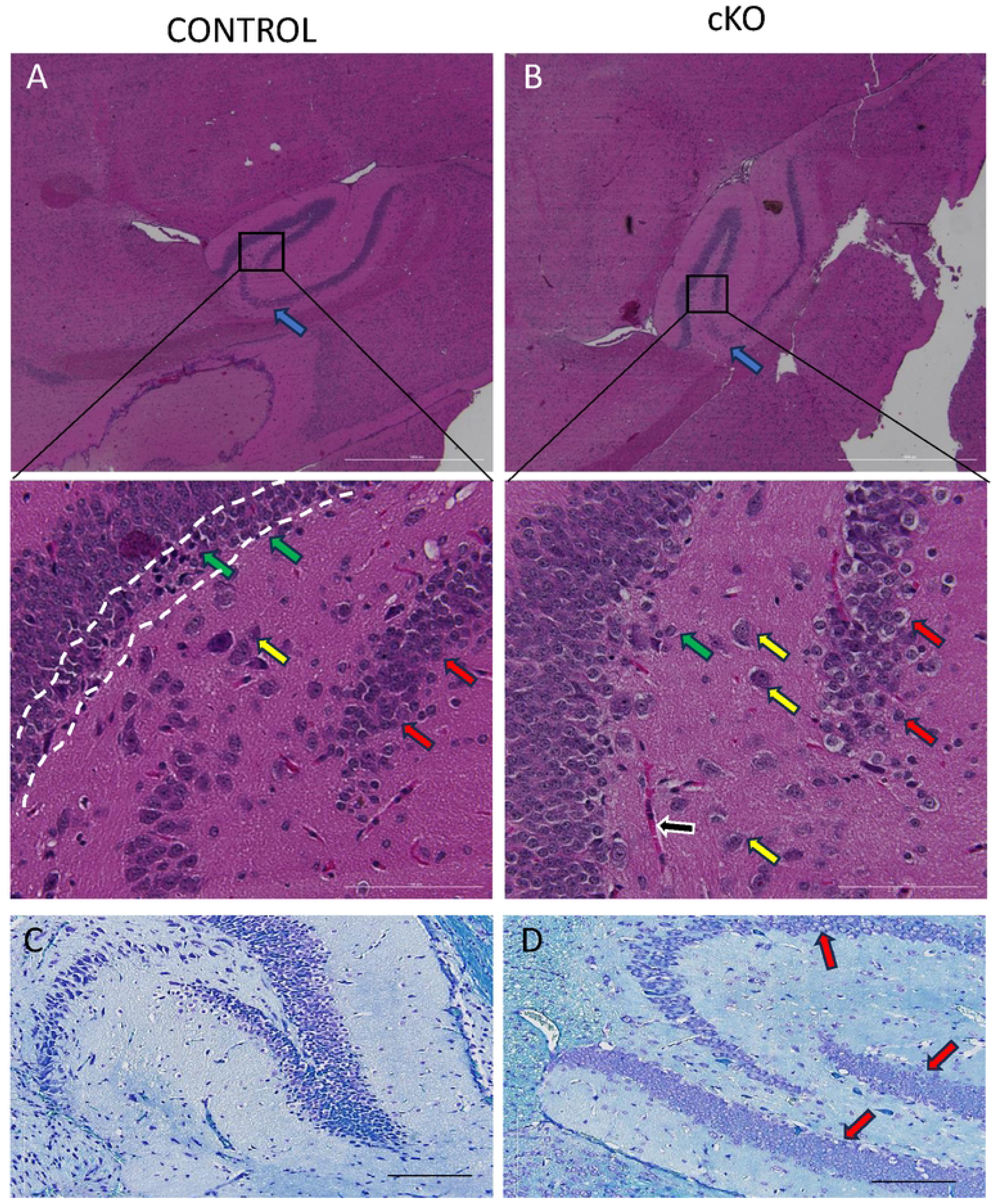
Hippocampus of the cKO mice have increased vacuolation of pyramidal neurons and a reduction of CA3 neurons and granular cells when compared to control mice. The polymorphic layer of dentate gyrus (A and B) with CA3 neurons are identified with blue arrows, the dense immature granules with green arrows. The white dotted line showing border of the dense immature granular layer, and yellow arrows identify pyramidal neurons of polymorphic layer, and red arrows identify granular cells. A congested capillary is present in the cKO hippocampus (black arrow). Hematoxylin and eosin staining of the hippocampus in A-B, and E-F is Luxol Blue staining. A and C are control mice, B and D are cKO mice. Scale bar = 1000 microns in A and B, 200 microns in the insets, and 100 microns in C and D.

Finally, the cerebellum of GABBR2 cKO mice demonstrated alterations in the organization of the Purkinje cell layer, with fewer Purkinje neurons, and more swollen or vacuolated cells (Figure 5A versus 5B, yellow arrows). Additionally, there were fewer dendritic projections penetrating into the molecular layer and reduced complexity of the dendritic branching pattern. (Figure 5A versus Figure 5B, red arrows).

**Figure 5.**
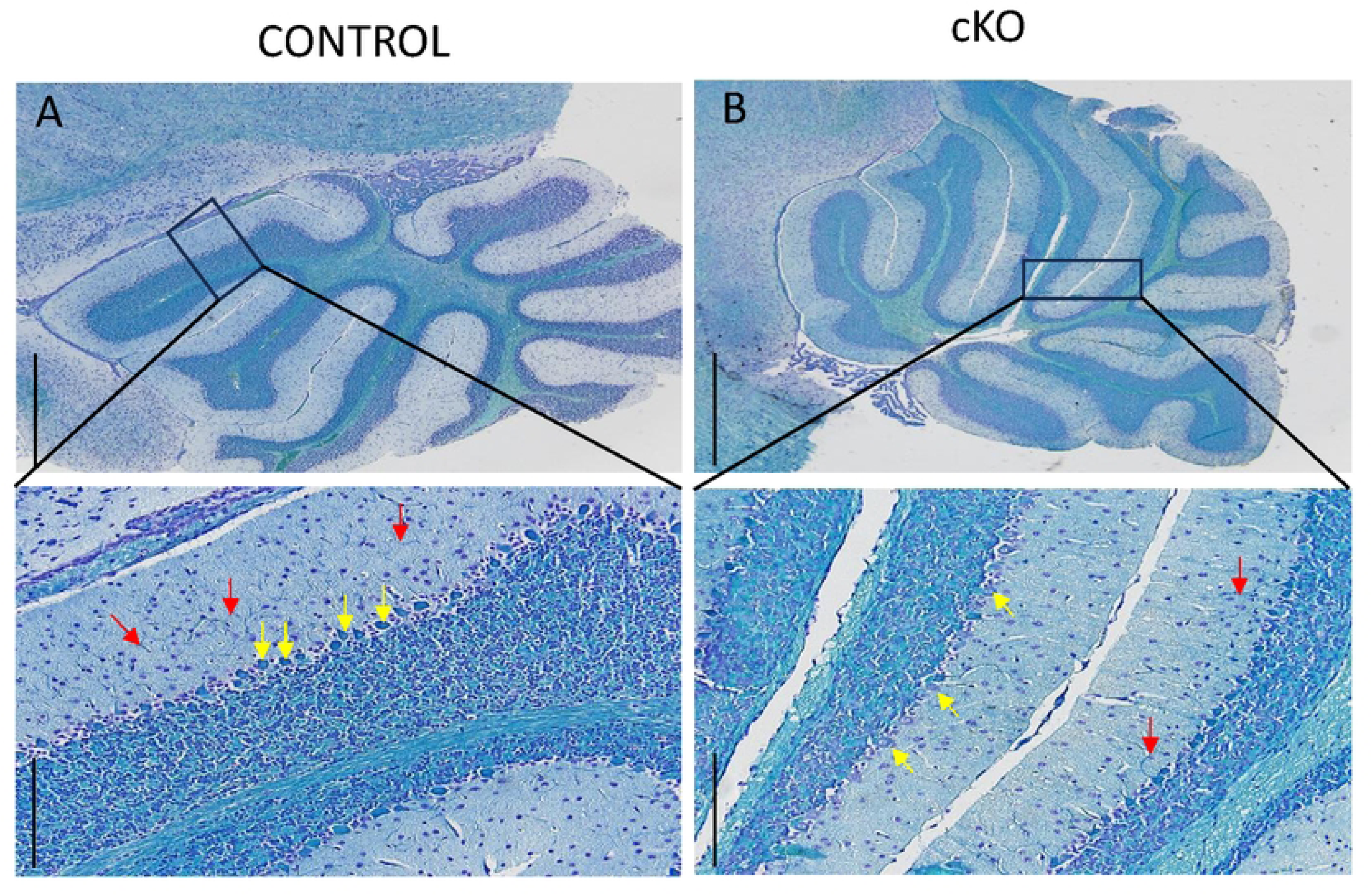
Cerebellum of cKO and control mice stained with Luxol blue. A is a control brain and B is a representative cKO brain. The cerebellum has reduced Luxol blue staining in the cKO compared to the control brain. There are fewer Purkinje neurons (yellow arrows) and shorter, fewer and less complex dendritic extensions (red arrows) in the cKO cerebellum (A versus B). A and B Scale bar is 1000 microns. Inserts of A and B scale bar is 200 microns.

### Loss of GABBR2 Alters Behavior and Motor Skills

Previous reports on the global GABBR2 KO mice described hyperalgesia, hyperlocomotion, elevated anxiety-related behaviors, and spontaneous seizure activity (3, 26). Therefore, we examined these activities in GABBR2 cKO mice to determine whether they mimicked the phenotype of global GABBR2 KO Mice. GABBR2 cKO mice demonstrated a greater than 3-fold increase in locomotor activity compared to control mice (Figure 6). There was a significant reduction of unsupported rearing behavior in GABBR2 cKO mice as compared to controls (Figure 6). This decrease in exploratory behavior is likely indicative of increased levels of stress but can also be seen in the setting of neurodegenerative disorders (27–29).

**Figure 6.**
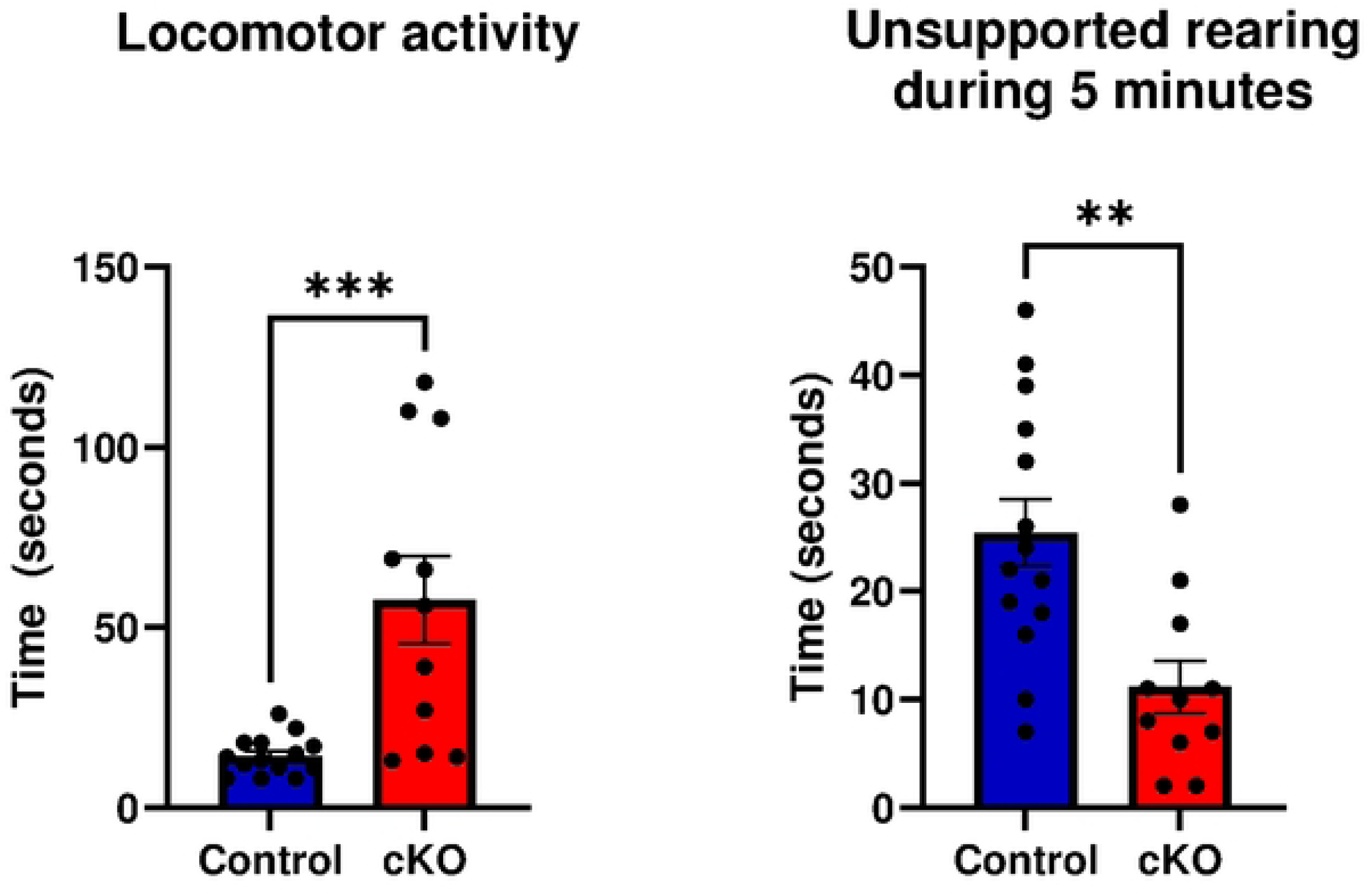
Measurements of locomotor activity and unsupported rearing in cKO versus control mice. Increased locomotor activity and decreased unsupported rearing is seen in cKO mice. Bars represent the mean ± SEM. ***p<0.001, **p<0.01. n= 14 for control mice, n=11 for cKO mice.

Baclofen is an agonist for gamma-aminobutyric acid (GABA) B receptors, and acts as a muscle relaxant (30, 31). Global GABBR2 knockout mice were previously shown to be refractory to baclofen as measured by changes in rotarod performance (3). Therefore, we assessed rotarod performance and responses to baclofen in GABBR2 cKO and control mice. During the rotarod training period preceding baclofen administration, it was clear that cKO mice of both sexes had a baseline decrease in their ability to remain on the rotarod (Figure 7A). Therefore, we expressed the response to baclofen as the change from baseline. Control mice of both sexes had a clear decrease in rotarod performance after baclofen treatment. However, despite the reduced performance at baseline, cKO mice showed no additional decline in performance after administration of baclofen (Figure 7B).

**Figure 7.**
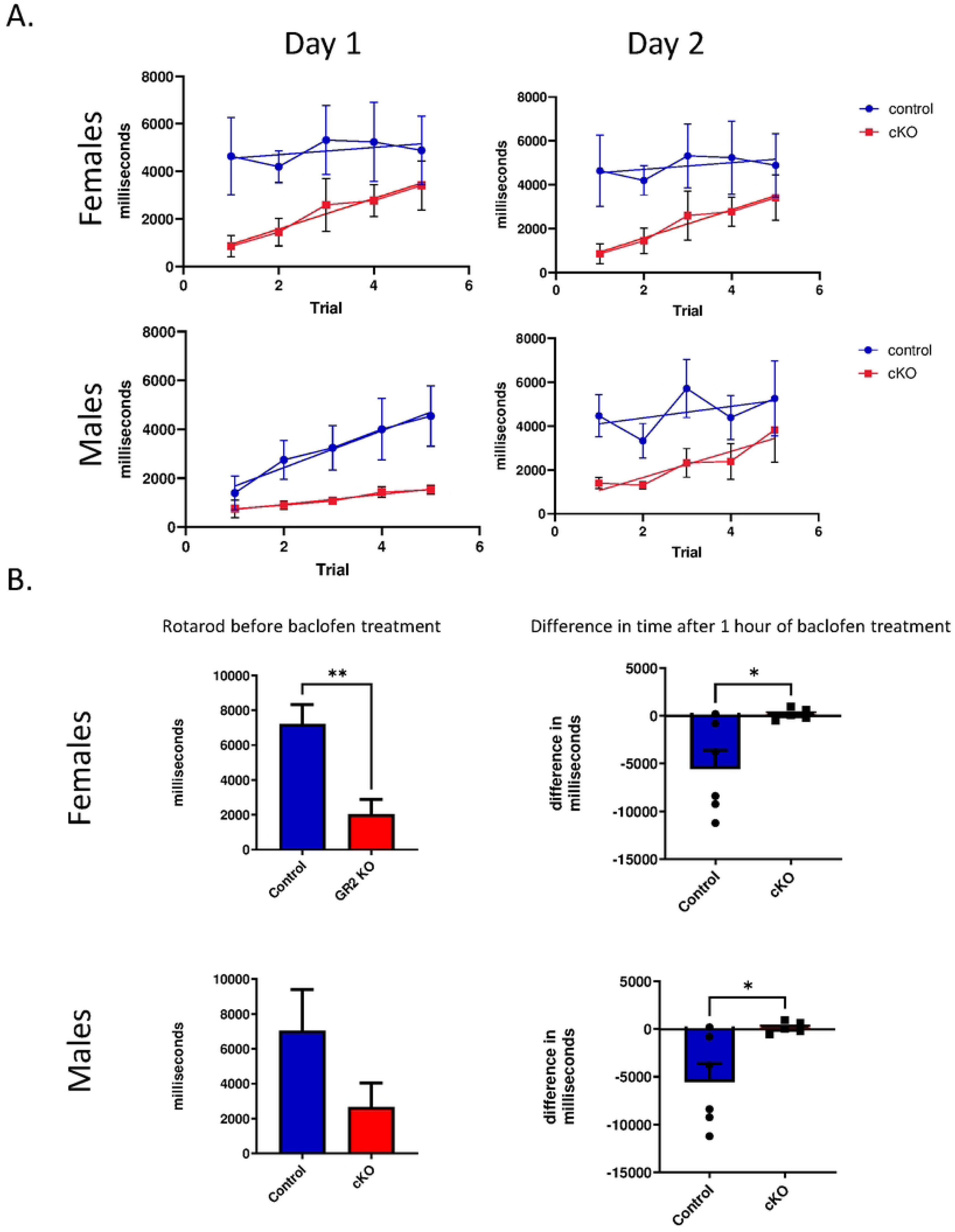
Rotarod experiments in female and male control and cKO mice. A). Training periods over 2 separate days showing times on rotarod for control (blue) and cKO mice (red). Points represent the mean ± SEM dwelling times over 5 trials on each of 2 days. Note that cKO mice stay on the rod for shorter periods at baseline, although they do improve with training. B). Rotarod experiments in female and male control and cKO mice before and after administration of baclofen, a GABBR2 agonist. Bar graphs on the left demonstrate the absolute dwell times on the rotarod. Bar graphs on the right represent the change in dwell time in response to baclofen administration. Bars represent mean ± SEM. *P<0.05, **p<0.01, ***p<0.001. n=4/group.

### Loss of GABBR2 results in seizures and premature death

We did not detect obvious spontaneous seizure activity while GABBR2 cKO mice were being monitored for locomotor activity. However, these mice had frequent seizures when subjected to stressful stimuli, such as, being handled or placed on the rotarod (Supplemental Video). In addition, we noted an increase in premature mortality in GABBR2 cKO mice. As shown in Figure 8, 100% of GABBR2 cKO mice died by 115 days of age while no control mice died during the same period. In addition to the pathological brain findings described above, necropsy of GABBR2 cKO mice showed little food in the stomach and small intestines, but no gross pathological changes. However, there was marked thymic necrosis. The spleen was enlarged and had areas of lymphocytic necrosis. The pancreas showed an absence of eosinophilic zymogen granules within the exocrine pancreatic acinar cells. Mice that had died some time before necropsy had brain findings similar to those described above. There was mild dilation of the lateral ventricles, multiple foci where there was decreased staining of the neuropil, especially in areas where cell nuclei were shrunken and there was cytoplasmic vacuolar degeneration. Diffuse demyelination was seen, blood vessels appeared congested, and some vessels contained mature fibrin.

**Figure 8.**
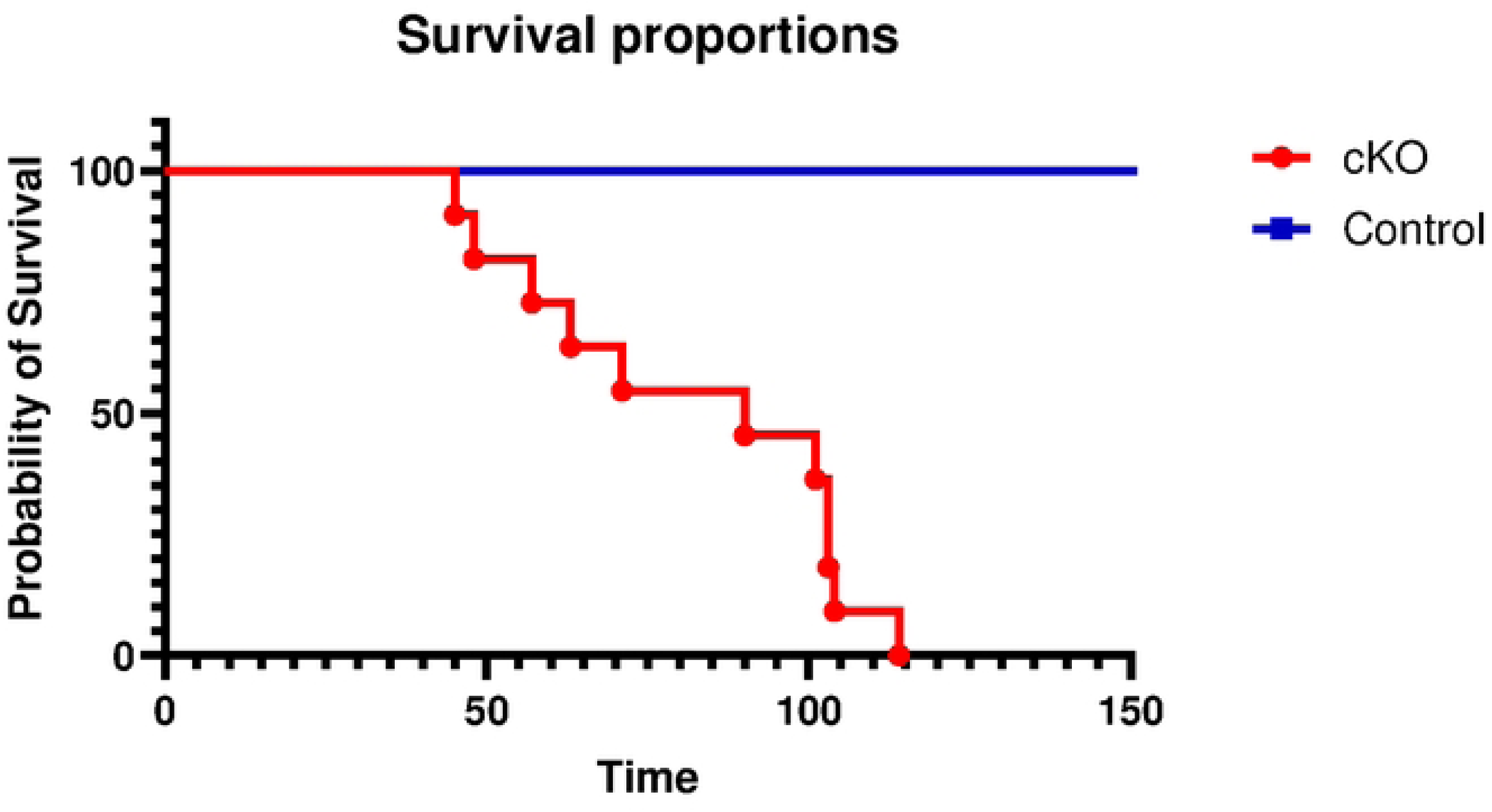
Kaplan-Myer plot showing survival curves of control and cKO mice. No control mice died over 150 days of observation. 100% of cKO ^mice^ died by 115 days. n= 11 in each group. Gehan-Breslow-Wilcoxon test (p<0.0001).

## Discussion

We generated a floxed GABBR2 mouse, that when crossed with Cre-expressing mice can be used to generate conditional KO mice targeting GABBR2 expression in different cell types. In this report, we crossed floxed GABBR2 mice with Actin-Cre mice to produce a widespread knockout of the *Gabbr2* gene. *Gabbr2* mRNA expression was reduced by over 80% in the whole brain and GABBR2 protein was not detected by immunoblot, documenting efficient disruption of the *Gabbr2* gene. The phenotype of Actin-Cre GABBR2 cKO mice was similar to the global GABBR2 KO mouse (3). These mice demonstrated increased locomotor activity but a decrease in unsupported rearing behavior. These mice also have impaired motor coordination and balance as measured by decreased ability to remain on a rotarod and seizures in response to being handled. These neuro-behavioral changes were accompanied by widespread changes in brain histology and also a reduced lifespan, both speaking to the importance of GABBR2 signaling for overall brain health and, perhaps, whole-body physiology as well.

We found that the loss of GABBR2 led to histological changes in different areas of the brain. In the cortex, GABBR2 is expressed in many different neurons including GABAergic cortical interneurons (32) and inhibitory interneurons (33). Loss of GABBR2 would be expected to impair slow inhibitory Gaba signaling to the interneurons connecting different regions of the cortex, perhaps resulting in a progressive decline in interneuron function. Loss of inhibitory interneuron signaling may also result in changes in cell viability as reflected here as a reduction of cortical thickness, a decrease in S100 staining, reductions in dendritic extensions and the presence of vacuolated neurons and cellular debris. Functionally, such a decline in neuronal populations might contribute to the loss of motor skills and cognitive function in the mice which we saw during the psychological and rotarod experiments (Figure 6 and Figure 7).

The hippocampus functions in memory and learning and GABBR2 expression typically occurs in the area near mossy fiber synapses that form the major excitatory input into the auto-associative network of pyramidal cells in the CA3 region (34). The loss of inhibitory input by GABBR2 to CA3 neurons could produce excitotoxity resulting of loss of pyramidal neurons (Figure 4B). Loss of neurons that govern lateral inhibition in the dentate gyrus can result in delamination of the granule cell layer and multilamellar discharges in response to cortical stimuli resulting in increased excitotoxicity (35). In the GABBR2 cKO mouse, progressive damage over time to the excitable neurons lacking GABBR2 input in the dentate gyrus of the hippocampus likely results in susceptibility to seizures and hyperexcitability (Figure 6). The histological phenotype of the GABBR2 cKO hippocampus is reminiscent of patients with epilepsy with a loss of dentate hilar neurons that govern dentate granule cell excitability (36, 37).

Cerebellar Purkinje neurons are known to express GABBR2 (38). Purkinje neurons project to the intermediate discharge layer and are the key efferent output of the cerebellum. The GABBR2 cKO mice have fewer Purkinje neurons (Figure 5, yellow arrows) and smaller dendritic arbors (Figure 5, red arrows) contributing to the abnormal rotarod performance. Purkinje dysfunction may also lead to fewer connections between the Purkinje neuron’s dendritic arbors and the interneurons in the molecular layer which may increase glutamatergic stimuli and neurotoxicity. Some seizure disorders cause increased Purkinje death by glutamate excitotoxicity (39).

In conclusion, the phenotype of GABBR2 cKO mice was similar to the global GABBR2 KO mouse (3). Our characterization of the Actin-Cre GABBR2 cKO mice has revealed histological changes in multiple areas of the brain in addition to changes in rearing behavior and premature death. Our studies do not discriminate between whether these changes are the result of altered brain development or due to progressive excitotoxic neuronal damage, although the use of inducible Cre transgenes could address this question in the future. Nevertheless, these studies demonstrate that the floxed mice reported here will provide a new tool to target tissue-specific GABBR2 signaling through Cre-mediated recombination. This will now provide scientists the ability to study GABBR2 function in different cell types without the potentially confounding effects of the neurological dysfunction caused by global knockout of GABBR2.

## Acknowledgments

We thank the Yale Genome Editing Center for their help in generating the GABBR2 cKO mouse.

## Notes

### Competing Interest Statement

The authors have declared no competing interest.

